# Single-shot detection of microscale tactile features

**DOI:** 10.1101/2024.09.03.610877

**Authors:** Sasha Reschechtko, Wylianne R. Pangan, Reza Zeinal Zadeh, J. Andrew Pruszynski

## Abstract

Tactile detection of very small features requires relative motion between the fingertip and a surface. The specific movement strategies that people use may be critical to maximize detection ability but little is known about the movement strategies people employ to support such detection. Here, human participants actively scanned a fingertip across a pair of silica wafers to detect which of the two contained a microscale feature (2, 6, and 10 μm height and 525 μm diameter). We constrained fingertip movement to ensure that participants would always contact the feature and would only contact the feature once. These procedures encouraged participants to use strategies that optimized detection rather than search and thus allowed us to more directly link movement strategies to detection. We also investigated the effects of fingertip movement direction and the finger used on detection. We found that participants were able to consistently detect microscale features as small as 2 μm on the basis of a single contact event. The contact forces that participants used were substantially higher than those observed in previous studies focused on tactile search or geometric feature extraction. Scanning speeds were slower than those found during tactile search but faster than those reported during geometric feature extraction. Taken in conjunction with the associations between detection and finger used as well as scan direction, our results suggest that control and consistency of fingertip movement may be a primary consideration for movement strategies that optimize tactile detection.

## Introduction

Whether moving our fingers over a roll of tape to find the edge or across a polished surface to feel for imperfections, we often sense features that are too small to be felt without relative motion (O’Connor et al., 2021) and may be difficult to see with the unaided eye. On very smooth surfaces, people are able to accurately detect circular features with a diameter of ∼600 μm and a height of ∼1.3 μm and edges ∼0.85 μm tall (Johansson & LaMotte, 1983). Although this detection ability is supported in part by the dense innervation of the human fingertips (Johansson & Vallbo, 1979), the presence and parameters of movement are also critical for detection (Hollins & Risner, 2000; Khamis et al., 2021; Manfredi et al., 2014). One example of the importance of movement in such circumstances comes from a study in which participants were asked to move their fingers to detect features with μm-scale heights (Kennedy et al., 2011). Although outright detection thresholds were minimally affected by age, the amount of time it took individuals to complete the detection task increased dramatically, pointing toward older individuals using different movement strategies or needing additional bouts of contact to extract the appropriate information.

While movement is required for tactile detection of small features, little is known about the active movement strategies that people use specifically to *detect* – rather than search for – these features. Callier and colleagues investigated movement strategies used to assess various aspects of textures (Callier et al., 2015), and a number of groups have focused on the search strategies that people use to find a macroscopic features or extract geometric information from macroscopic features (Morash, 2016; Olczak et al., 2018; Smith et al., 2002; Vega-Bermudez et al., 1991) rather than strategies for detection itself. Previous studies that focused directly on the detection of microscale features (Johansson & LaMotte, 1983; Kennedy et al., 2011; LaMotte & Whitehouse, 1986) have imposed few constraints on movement and did not measure the detection behaviors people employed. In particular, these studies did not restrict participants to a single instance of contact between the finger and the feature, instead allowing participants to move their fingers across these features multiple times in a single prescribed direction (Johansson & LaMotte, 1983) or without constraints (Kennedy et al., 2011). As such, the specific movement parameters that are conducive to detecting microscale features remain unknown.

In the present study, we performed experiments in healthy young adults that quantified perceptual thresholds and the behavioral strategies used to detect microscale (2, 6, or 10 μm in height) features on the basis of a single contact event. We additionally investigated whether the finger used or direction of finger movement across the feature affected tactile detection accuracy and behavioral strategies. Our participants performed a two-alternative forced choice task to identify which of two surfaces contained a microscale feature with the direction of movement enforced by a physical aperture placed over the stimuli. The aperture guaranteed that participants would contact the feature when it was present so that participants did not have to search. Participants were allowed only a single finger sweep over each surface so that tactile detection had to be based on a single contact event. We report that participants are able to reliably identify the presence of microscopic features, even for the 2 μm feature height. This identification is more reliable when using the index finger than the little finger, and sweeps in the anterior-posterior direction (i.e. toward the body) are more successful than those in the medial-lateral (i.e. left-right) direction, but these advantages are only present for the smallest feature. We also report that participants use substantially larger contact forces in this context than previously reported in studies where participants extract geometric information about macroscale features or make texture judgements (Lederman, 1974; Olczak et al., 2018; Vega-Bermudez et al., 1991).

## Methods

### Participants

33 healthy individuals (age 19-36) participated in this study, which was broken into two identical experimental sessions, one using the index finger and one using the little finger. 27 of these individuals (18 women and 9 men; 7 left handed) participated in both experiments; 3 individuals (all right handed and female) participated in experiment 1 only and 3 other individuals (also all right handed and female) participated only in experiment 2. For those who performed both sessions, the order of fingers used was counterbalanced: 13 participants used their index finger first and 14 participants used their little finger first. Participants provided informed consent and performed experiments according to procedures approved by the Institutional Review Board of the Human Research Protection Plan at San Diego State University.

### Tactile stimuli

The Western Nanofabrication Facility (London, ON, Canada) manufactured our stimuli. Each stimulus was produced by depositing a gold mask with a chromium adhesion layer through a 525 μm diameter aperture onto the center of a 100mm fused silica wafer. The wafer was then etched in buffered hydrofluoric acid (etch rate of 0.1 μm/min) to obtain the desired feature height and bonded to a 17-7PH stainless steel backing plate for protection. The height of the etched features was verified using a stylus profilometer (Tencor P7, KLA Corporation, Milpitas, CA, USA).

### Procedure

Before starting the experimental trials, participants washed their hands with soap and water and dried them. We did not control the room environment; average room temperature was 18.8°C (range: 18.4-23.4) and average humidity was 65% (range: 44-85%).

During each experimental trial, we presented participants with two silica wafers: one had a feature (2, 6, or 10 μm height; 525 μm diameter) and the other had no feature (i.e. was “blank”). A 3D-printed aperture covered most of the wafer surface except for an open slot (18×85 mm) that constrained the participant’s finger motion to the medial-lateral or anterior-posterior direction. We instructed participants to feel both wafers in succession using the pad of the index or little finger (depending on the experiment) of their dominant hand and then indicate which wafer had the feature on it by pressing a microswitch corresponding to that wafer. We did not prescribe the order in which participants touched the wafers. We instructed participants to feel each wafer once per trial in a single sliding motion, so they could not reverse directions of movement or move their finger back and forth on the wafer. Between trials, the experimenter switched wafers according to a quasi-randomly generated sequence (including 3 wafers with features and 5 blank wafers) to ensure each participant felt each feature the same number of times. If the sequence called for the same feature to be presented in the same position twice in a row, the experimenter removed the wafers from the apparatus before re-inserting them. A blind blocked the participant’s view of the wafers so that they could not keep track of which wafers were being presented. Each wafer with a feature was presented 23 times per sweep direction, for a total of 138 trials per experiment (23 trials per height × 3 heights × 2 directions).

Experimental flow was controlled and data were logged using a program written in the LabVIEW programming environment (NI, Austin TX, USA). We recorded contact kinetics at 500 Hz using a 6-axis force/torque transducer (Axia80-M8, ATI Industrial Automation, Apex NC USA; force resolution: 0.04 N, torque resolution: 0.002 Nm). The transducer was placed under the fixture supporting both wafers. Because participants contacted each wafer in succession using a single finger, the forces and torques measured by this transducer could be used to compute the center of pressure (COP) of force applied by the fingertip, which we considered to be fingertip position. We computed the position of the fingertip on the anterior-posterior/medial-lateral plane as: [X, Y] = [τ_x_ / F_z_, τ_y_ / F_z_], where τ represents torque and F represents force about the subscript axis. Participants indicated which wafer they thought had the feature on it by pressing one of two microswitches corresponding to their choice; this keypress was logged and used as a signal to stop data collection for the trial. Participants did not receive any feedback about whether their selections were correct during or after the study, although the experimenter asked how confident they felt in their performance after the visit. An illustration of the apparatus and coordinate system, as well as the microscale features, is presented in Figure 1.

**Fig. 1.**
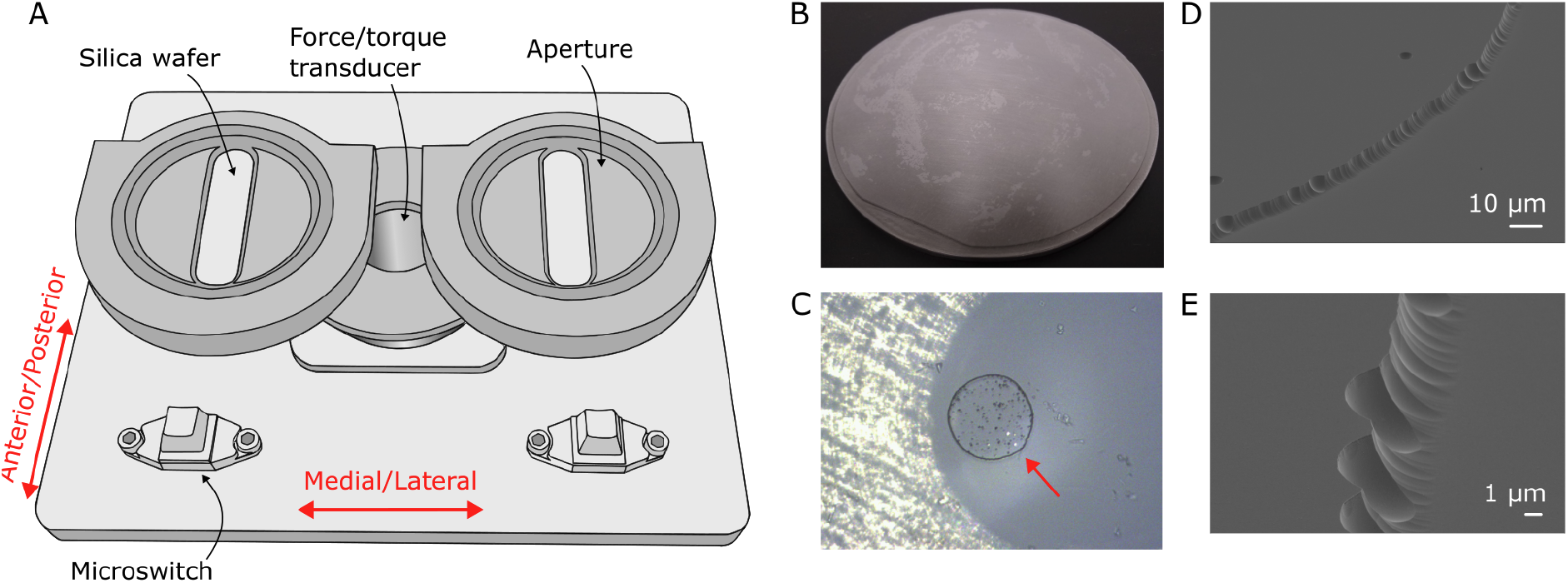
Experimental Setup. A: Apparatus used to hold wafers during experimental trials. Two silica wafers were loaded into the apparatus; only the area under the aperture was exposed to participants. Participants used the corresponding microswitch to indicate which wafer they thought had a microscale feature. Directions of movement (Anterior/Posterior and Medial/Lateral) shown with red arrows. In the illustration, the apertures are oriented for Anterior/Posterior sweeps. B: Image of a silica wafer mounted on its protective steel backing plate. The feature is not visible to the naked eye; the visible asperities are in the adhesive layer between the wafer and the mounting plate. C: Photograph under microscopy of a feature seen from above; red arrow indicates the perimeter. Visible asperities are due to the adhesive layer between the wafer and the backing plate and machining marks on the backing plate. D: The edge of an exemplar 4 μm feature manufactured using the same process as features used in the present study under scanning electron microscopy; the scale bar is 10 μm. E: The edge of a 4 μm feature under higher magnification scanning electron microscopy; the scale bar is 1 μm.

### Analysis

We analyzed each trial to find the time when the participant’s finger was in contact with the wafer to identify times of interest for analyzing contact forces. Due to the two-alternative forced-choice design, each data file contained recordings of two contact events (one per wafer). We identified onset and offset times for each contact using the normal force participants applied to each wafer; we defined contact onset as the time when normal force reached a trial-specific threshold of 0.05 N greater than the average value over the first 200 ms of the trial. We investigated a variety of threshold values and they minimally influenced the computed contact times. In cases where there were more than two epochs when normal force exceeded the threshold (e.g. participants bumped into the fixture between wafers), we identified the relevant contact events as the two longest events. To analyze movement speed, we computed the center of pressure of the fingertip force applied for each contact event and computed the derivative of the center of pressure timeseries. For each participant, we removed trial values of speed and contact forces that were > 3 SD from the average observed during each finger × direction condition, as such values likely indicated erroneous contact identification. At most, this criteria removed two trials per participant from analysis (out of 68 trials completed).

### Statistics

Statistical analyses were performed using JASP 0.19.0 (JASP Team, 2024). We quantified tactile detection performance as the proportion of correct responses. Based on a binomial test with size parameter 23, successful detection of the location of the stimulus at least 16 times (proportion of correct responses ≥ 0.7) corresponds to a 5% or lower likelihood of detection by chance. We therefore used this cutoff to evaluate performance against chance levels.

We analyzed the effect of finger (Index and Little), movement direction (anterior-posterior and medial-lateral), and feature height (2, 6, 10 μm) on proportion of correct responses using linear mixed effects models with Finger, Movement Direction, and Feature height as fixed effects and participant as a random effect. When investigating speeds and contact forces, we did not investigate the effect of feature height because participants could not predict feature height – so there was no causal basis for interpreting any differences in behavior as a result of feature height. For linear mixed effects models, degrees of freedom and corresponding p-values were estimated using the Satterthwaite method (per JASP default). We used Spearman rank-order correlation to investigate potential relationships between success rates and applied force, speed, and fingertip area. Unless otherwise noted, descriptive statistics are reported as mean ± standard deviation.

## Results

Participants took approximately one hour to complete each experimental session (one per finger used). 27 participants took part in both experimental visits, while 6 participants completed only one visit (3 index finger only; 3 little finger only) such that 30 participants’ data were analyzed for both finger conditions. During each visit, participants performed 138 trials (23 per feature height per direction). Participants’ self-reported confidence in their performance ranged widely.

### Psychophysics

Participants’ performance was highly influenced by the height of the feature they felt and finger used but on average it was above chance for all feature heights. For the index finger, 77% of participants performed above chance in anterior-posterior sweeps and 57% performed above chance in medial-lateral sweeps. For the little finger, 60% of participants performed above chance for anterior-posterior sweeps although only 23% of participants performed above chance for medial-lateral sweeps.

Across participants, sweep direction, and finger, the average proportion of correct identifications was 0.72 ± 0.14 for 2 μm features, 0.93 ± 0.10 for 6 μm features, and 0.96 ± 0.08 for 10 μm features. The proportion of correct identifications for the index finger was higher on average than for the little finger (0.89 ± 0.15 versus 0.85 ± 0.16; linear mixed effects model factor *Finger*: F_2,30.12_; *P* = 0.002). Correct identifications were also higher for anterior-posterior direction of finger sweep than for medial-lateral sweep direction (0.89 ± 0.14 versus 0.85 ± 0.16), reflected in a significant interaction between *Feature Height* and *Direction* (F_2,30.32_; *P* = 0.01). Individual contrasts for *Feature Height* and *Direction* interaction revealed that the sweep direction significantly affected detection of features with lower heights (2 and 6 μm), but not the tallest feature. Although the interaction between *Finger* and *Feature Height* was equivocal (F_2,29.86_; *P* = 0.08), a similar pattern was seen in the individual contrasts: index finger performance was better for the 2 μm feature only. These results are shown in Figure 2A-B. For the index finger, participants’ success in the task during sweeps in one direction was associated with their success using sweeps in the other direction (Spearman correlation: ρ = 0.671; P = 4.867e^-5^), but this association was not present for the little finger (ρ = 0.218; *P* = 0.246). These results are shown in Figure 2C.

**Figure 2.**
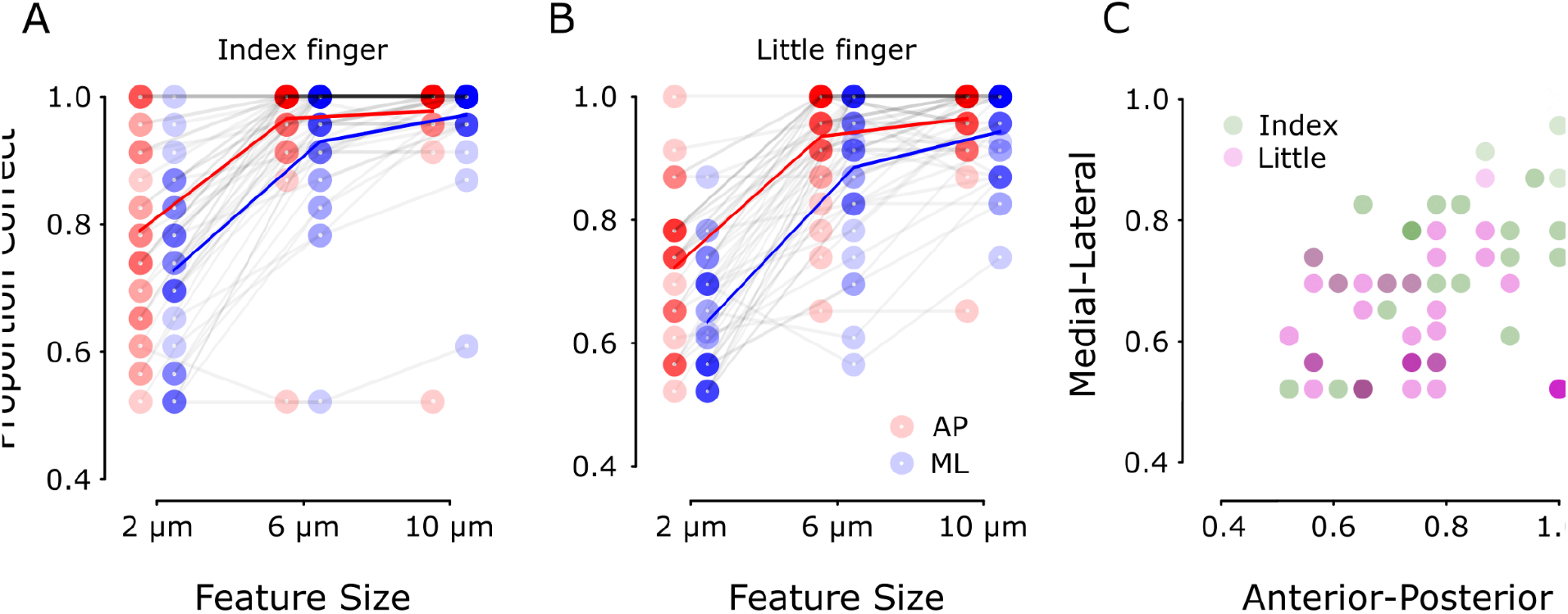
Psychophysics results. Panel A: Proportion of correct responses that participants made when using their index fingers to detect features of denoted heights in a two-alternative forced-choice task. Panel B: Proportion of correct responses when participants used their little fingers for the detection task. Each dot represents one participant (n = 30) in one finger x feature height x sweep direction condition. Red denotes anterior-posterior and blue medial-lateral sweeping movements. Red and blue lines indicate across-participant average performance for anterior-posterior and medial-lateral sweeps, respectively. Panel C: Proportion of correct responses in the 2 um condition for anterior-posterior sweeps versus medial-lateral sweeps.

### Contact forces

Participants deployed larger normal forces with the index finger (2.25 ± 1.84 N) than the little finger (1.77 ± 1.20 N). The linear mixed effects model indicated this difference was significant (factor *Finger:* F_1,29.26_ = 5.91; *P* = 0.021), but we did not observe significant effects of the sweep *Direction* (F_1,50.92_ = 1.699; *P* = 0.198) or a significant *Finger* × *Direction* interaction (F_1,57.67_ = 1.80; *P* = 0.185). These results are shown in Figure 3A. The normal forces that participants deployed were strongly correlated across fingers and directions of finger sweep: individuals who applied more force in a given *Finger* × *Direction* combination tended to apply more force in all other combinations as well (Spearman correlations: all ρ > 0.75; all *P* < 1.0e^-6^). These results are shown in Figure 3C.

**Figure 3.**
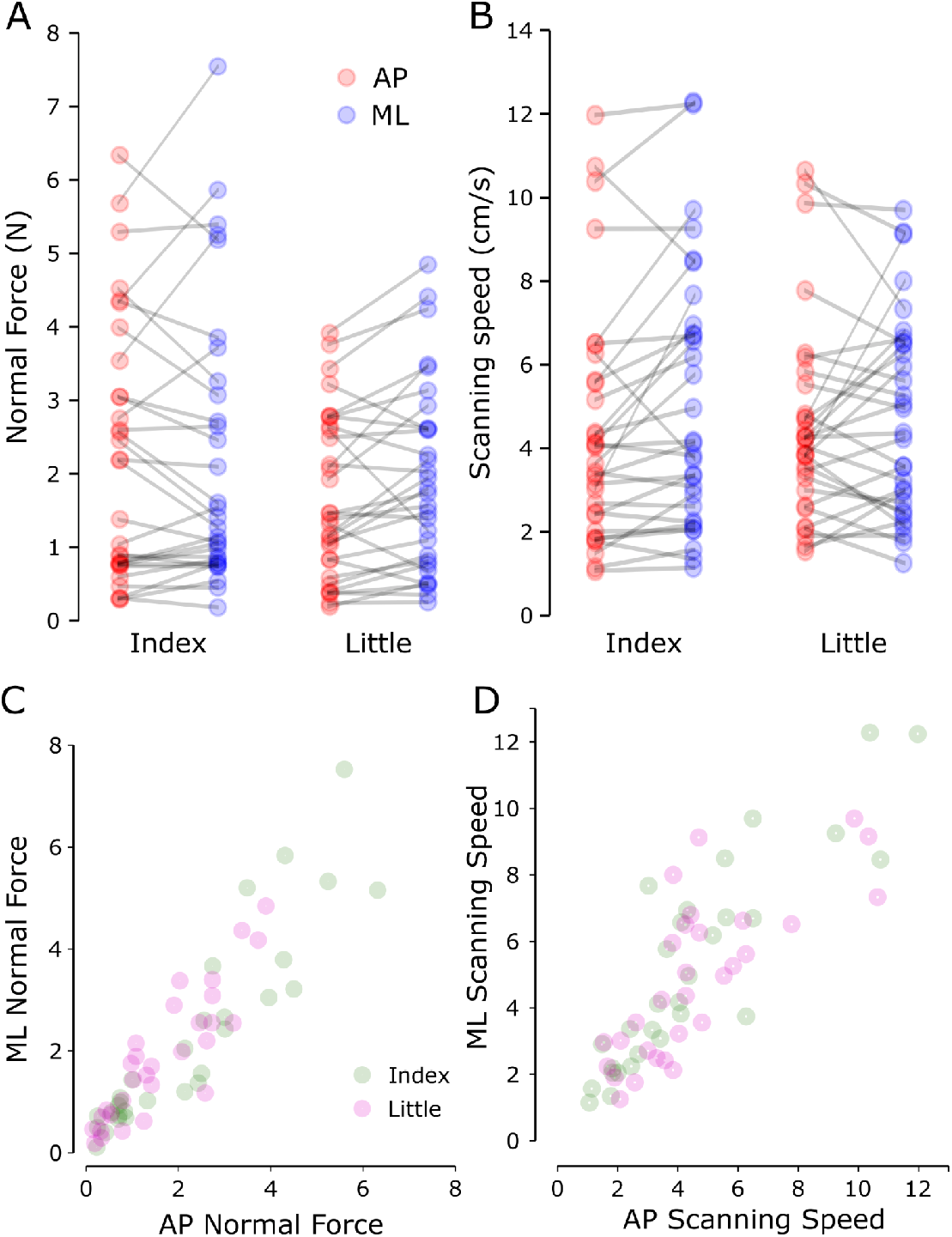
Kinetics and Kinematics of tactile detection. Panel A: Normal forces deployed at the fingertip during tactile detection tasks. Panel B: Fingertip sweep speed during the tactile detection tasks. For panels A and B, Anterior-posterior sweeps are denoted in red and medial-lateral sweeps in blue. Panels C,D: scatterplots of normal forces and scanning speeds for anterior-posterior sweeps versus medial-lateral sweeps. For all panels, values are averages across all contact events and all trials for a given finger × direction combination. Each dot represents one participant (n = 30) in one finger × direction combination across all feature heights (69 trials less any discarded for technical reasons). Index finger sweeps are denoted in green and little finger in magenta.

For the index finger, normal force applied was significantly correlated with success in anterior-posterior sweeps (ρ = 0.488; *P* = 0.006) and medial-lateral sweeps (ρ = 0.436; *P* = 0.016). We did not find a reliable correlation between normal force and acuity for the little finger (anterior-posterior sweeps: ρ = -.026; *P* = 0.892; medial-lateral sweeps: ρ = 0.215; *P* = 0.235). These results are shown in Figure 4A (for index finger) and 4C (for little finger).

**Fig 4.**
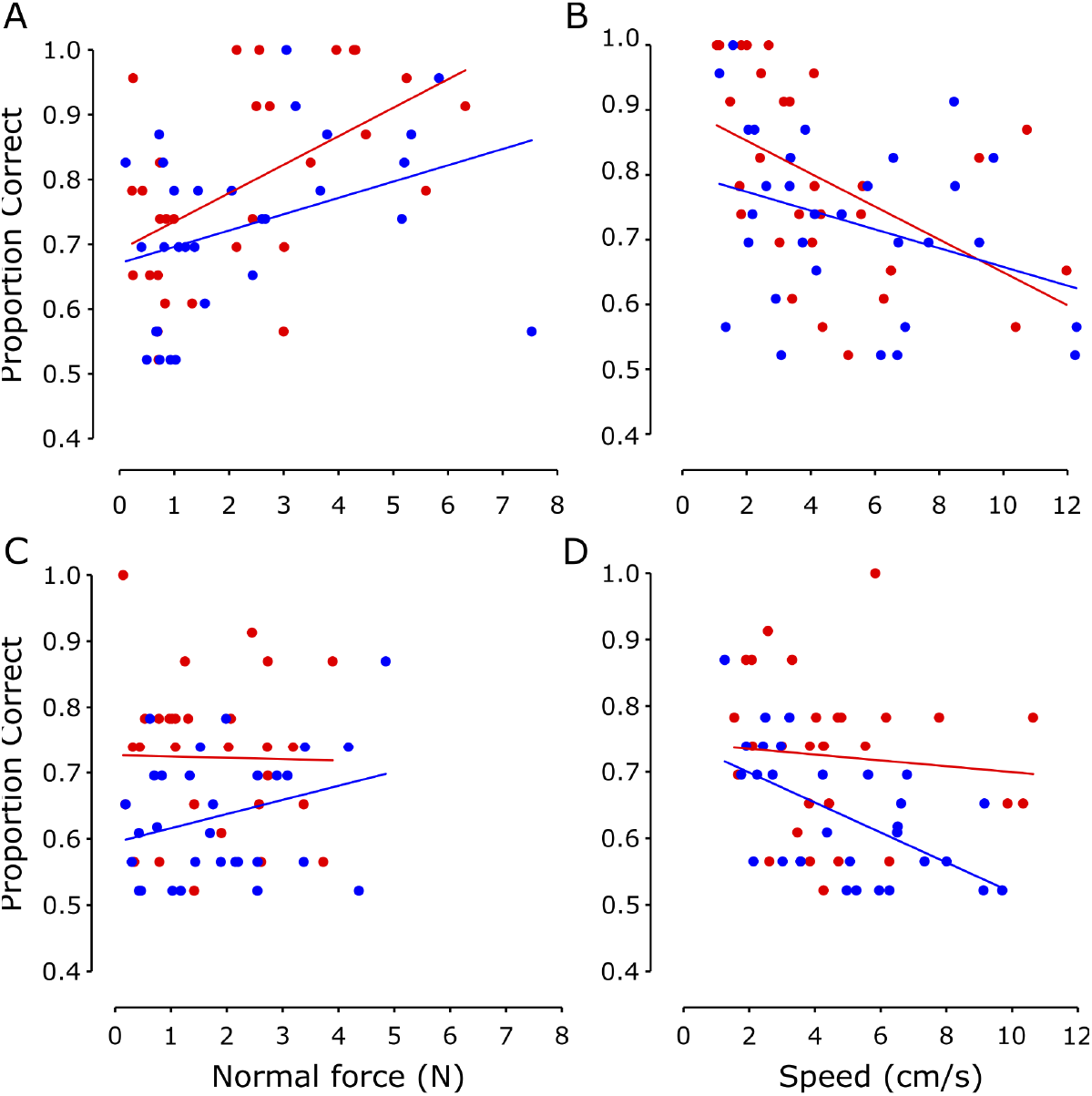
Kinetic/kinematic variables and tactile detection success. Top row: Index finger; bottom row: little finger. Panels A and D: average normal force deployed during the 2 μm detection tasks and success rates in identifying which wafer had the feature (proportion correct).Panels B and E: average tangential force deployed in the direction of movement during the 2 μm detection tasks and success rates in identifying which wafer had the feature (proportion correct). Panels C and F: average scanning speeds used the 2 μm detection tasks and success rates in identifying which wafer had the feature (proportion correct). For all figures, each dot represents a single participant (n = 30) in a single finger x direction condition. Red dots indicate anterior-posterior sweeps; blue dots indicate medial-lateral sweeps.Lines indicate best-fit regressions.

### Scanning speeds

We computed the speed that each participant used to move their finger across the silica wafers by calculating center-of-pressure as a function of time (see Methods). Because participants could not predict which feature height they would be exposed to on a given trial, we investigated the effects of movement direction and finger on fingertip speeds across all feature heights.

Across participants, fingers, and movement direction, average velocities were variable and only appeared sensitive to movement direction: a linear mixed effects model with fixed factors *Direction* (2 levels) and *Finger* (2 levels), revealed a marginal effect of Direction (F_1,31.58_ = 4.484; *P* = 0.042), with medial-lateral sweep velocity higher on average (5.0 ± 2.8 cm/s) than anterior-posterior (4.5 ± 2.7 cm/s). We did not observe a significant effect of the finger used (F_1,28.35_ = 0.100; *p* = 0.754) or a *Finger* × *Direction* interaction (F_1,29.94_ = 1.742; *P* = 0.197). The average speed was very similar for index finger and little finger (4.8 ± 3.0 versus 4.7 ± 2.4 cm/s). These results are illustrated in Figure 3B.

Participants who moved their fingers faster in one direction tended to move their finger faster in the other direction as well. For the index finger, higher AP speed was associated with higher ML speed (ρ = 0.887; *P* = 3.713^-7^), and we observed the same relationship for the little finger (ρ = 0.720; *P* = 3.689e^-5^). Similarly, participants who moved their index finger faster tended to move their little finger faster (AP: ρ = 0.709; *P* = 5.546e^-5^, ML: ρ = 0.838; *P* = 1.665e^-6^). These results are shown in Figure 3D. Success for the index finger was significantly correlated with sweep speed in the anterior-posterior direction (ρ = -.625; *P* = 2.246e^-4^ but the association was equivocal in the medial-lateral direction (ρ = -.307; *P* = 0.098). In contrast, little finger success was not significantly correlated with sweep speed in anterior-posterior direction (ρ = -0.114; P = 0.550) but there was a significant correlation between speed and success in the medial-lateral direction (ρ = -.560; *P* = 0.001). These results are illustrated in Figure 4B (for index finger) and 4D (for little finger).

## Discussion

Participants in our study detected the presence of microscale features on one of two otherwise smooth silica wafers. Critically, they only made a single sweep of each wafer and were constrained such that they were guaranteed to contact the feature. Due to this constraint and trial structure, participants did not have to search for the feature but rather had to tune their behavioral strategies to optimize detection from a single contact event.

We found that single-shot detection was possible and detection thresholds were consistent with previous studies that allowed multiple contact events (Johansson & LaMotte, 1983; Kennedy et al., 2011). Even when participants were given only one chance per trial to feel a microscale feature, 23 of 30 participants could detect the shortest feature we used – 2 μm height – at above-chance levels with their index finger in the anterior-posterior sweep direction. More than half of participants performed above chance in all conditions other than medial-lateral sweeps with the little finger (where only 7/30 participants performed above chance). For the 2 μm feature height, participants performed better when using the index finger than the little finger, and they performed better when sweeping their finger in the anterior-posterior direction (along the long axis of the finger) than the medial-lateral direction, although this difference was most pronounced for the little finger. For the taller feature heights (6 μm and 10 μm), both fingers and sweep directions were almost uniformly successful and average success rates were in excess of 90%.

### Participants used relatively large forces to detect very small features

The features in this study were very close to previously reported detection thresholds and we expected that being successful would require finesse in terms of the forces they applied to the wafers. However, the forces our participants deployed were about one order of magnitude greater than those described in previous research – including studies detecting much larger features (Lederman, 1974; Olczak et al., 2018; Smith et al., 2002; Vega-Bermudez et al., 1991). Studies in which participants actively moved their fingers to feel macroscopic features have also reported lower levels of normal force application. Olczak and colleagues (2018) reported average forces of less than 0.5 N (and a range of approximately 0.1 to 0.7 N) in a study where participants moved their fingers across a series of 5 mm tall edges to determine their relative orientation. Smith and colleagues (2002) investigated the forces participants used to search for features of various sizes that were 620 μm tall or recessed; those individuals also deployed forces around 0.5 N.

Previous studies using stimuli similar to the ones here have shown that humans are able to detect similar microscale features when they actively move their fingertip across the feature as when the feature is passively scanned across the fingertip (Johansson & LaMotte, 1983; LaMotte & Whitehouse, 1986). But to our knowledge no direct comparisons of forces exist. In their active touch paradigm, Johansson and LaMotte did not record the forces that participants used. In LaMotte and Whitehouse’s passive touch paradigm, features were scanned across the fingertip with forces of 0.2 N and lower on the basis of fingertip indentation (0.5 - 1.0 mm) rather than the resultant contact force. It is notable, however, that their participants were able to perform the task – indicating that the high forces used here are not required for discrimination of microscale features.

What explains the higher forces used in our study? One possibility is that increasing contact force increases the sensitivity of peripheral tactile neurons and so, when given the chance in our active touch paradigm, participants use higher forces and take advantage of this added sensitivity. However, peripheral recordings from from macaque monkeys with stimuli similar to our own found that changes in force did not affect the neural responses recorded from FA-1 neurons innervating Meissner corpuscles, which were the only peripheral tactile neurons active during contact with features of 6 μm or smaller (LaMotte and Whitehouse, 1986). Additionally, although we can not rule it out, it seems unlikely that the additional force recruited other types of peripheral tactile neurons. For example, FA-2 neurons innervating Pacinian corpuscles are incredibly sensitive (O’Connor et al., 2021; Saal & Bensmaia, 2014), but LaMotte and Whitehouse (1986) found that macaque FA-2 neurons were particularly insensitive to features like the ones used here: while they recorded from FA-2 neurons that had very low force thresholds, those units did not respond to contact with features with heights under 21 μm.

Perhaps the more likely reason our participants deployed higher forces is to make their fingertip sweeps more predictable by minimizing other sources of tactile noise, in particular “stick-slip” events (Derler & Rotaru, 2013; Smith et al., 2002). When the fingertip moves along a very smooth surface, it has a tendency to stick and slip, which in the present study is much more salient than the feature itself. Therefore, moving the fingertip smoothly across the surface is much more important with microscopic stimuli compared to studies in which participants felt larger features. The coefficient of friction of the fingertip tends to *decrease* as contact pressure *increases* (Derler et al., 2009; Derler & Gerhardt, 2012) due to a variety of interrelated factors (reviewed in Derler & Gerhardt 2012). For this reason, higher levels of normal force may have made it easier for participants to minimize stick-slip events. This explanation could also be consistent with LaMotte and Whitehouse’s finding that humans can detect similar stimuli at low force levels during passive stimulation (LaMotte & Whitehouse, 1986). Passive stimulation applied via a motorized device likely reduced stick-slip events by moving the stimuli at a very consistent speed and force.

### Scanning speeds for single-shot detection

People seem to deploy fairly goal-specific movement strategies when performing various tasks related to the sense of touch. Callier and colleagues (2015) found that movement speeds varied depending on the property (roughness, slipperiness, or hardness) being assessed. They reported the highest average speeds (∼108 mm/s) for assessment of slipperiness, and lowest average speeds (∼49 mm/s) for assessing hardness, in addition to systematic changes in movement path depending on the surface characteristic to be assessed. Scanning speeds of 60-110 mm/s have been observed when participants perform tactile search tasks (Morash, 2016; Smith et al., 2002). In contrast, when participants moved their fingers to investigate geometric features of macroscale stimuli (Olczak et al., 2018; Vega-Bermudez et al., 1991), scanning speeds were lower at around 20 mm/s. Our participants used average scanning speeds of 45-50 mm/s, which may point to the specific nature of the detection task where search was unnecessary and participants did not need to remember and compare information across successive stimuli (Olczak et al., 2018) or compare those stimuli to known features like letters (Vega-Bermudez et al., 1991).

LaMotte and Whitehouse’s study indicates that detection of features with heights under 6 μm are likely to engage only FA-1 neurons, at least in the macaque monkey (LaMotte & Whitehouse, 1986). FA-1 neurons have extensive distal arborizations that enable a single neuron to innervate many mechanoreceptive end organs (Cauna, 1956, 1959; Nolano et al., 2003), resulting in spatially complex receptive fields that span multiple fingerprint ridges (Jarocka et al., 2021; Pruszynski & Johansson, 2014; Suresh et al., 2016; Vallbo & Johansson, 1984). Tactile detection of a microscale feature likely results from the temporal patterns of activity both within and across neurons (Hay & Pruszynski, 2020; Pruszynski et al., 2018) that are evoked by the lateral deflection of fingerprint ridges (Johansson & LaMotte, 1983; LaMotte & Whitehouse, 1986). Given the geometry of FA-1 receptive fields, a sweep of the fingertip across a microscale feature will likely result in a complex pattern of action potentials, as it does for macroscale features (Pruszynski & Johansson, 2014; Sukumar et al., 2022), and could help distinguish the sensations related to contacting a feature from those related to fingertip strains as the fingertip traverses the glass wafer. Our participants displayed average scanning speeds that are consistent with a recent report showing that human FA-1 neurons provide maximum information for discriminating between macroscale geometric features based on the patterning of their responses for scanning speeds at or below ∼50 mm/s (Sukumar et al., 2022). Although its difficult to compare these findings to our study given that our stimulus and task required detection rather than discrimination, it may be that participants tuned their scanning speed to move as quickly as possible – to limit stick slip events and to reduce overall time in the experiment – but not above the point where FA-1 neurons start to lose their capacity to signal the requisite information.

### Better performance for certain scanning speeds and fingers

In an active touch situation where participants move their finger volitionally to support tactile detection, participants could use information about how they intended to move their finger and surface conditions to generate a “forward model” (Miall & Wolpert, 1996; Wolpert et al., 1995) of the expected sensory experience associated with scanning a featureless surface. Deviations in sensory inputs from that expectation (due to the complex response to multiple fingerprint ridges contacting the feature in sequence) could be attributed to detecting the feature itself. In this case, an important strategy for detection would be making the sensory experience as predictable as possible. In addition to using higher forces to minimize stick-slip, the goal of predictability may explain the effect of scanning direction on performance, as our participants performed better using anterior-posterior sweeps to detect features. This may be due to participants having better control over the fingertip when sweeping in the anterior-posterior direction because all interphalangeal joints and a complex flexor/extensor network (Chao et al., 1976; Darling et al., 1994; Darling & Cole, 1990) can be engaged to help guide fingertip movement and tune compliance. In contrast, only one degree of freedom is available in the medial-lateral direction across the metacarpal phalangeal joint, and it is spanned by only a single muscle pair. Controllability may also be a factor explaining why participants performed better with their index fingers than little fingers, especially in the medial-lateral direction where 17 participants were successful using their index fingers to detect a 2 μm feature but only 7 were successful using their little finger. The additional controllability of the index finger could at least partially compensate for reduced controllability in the medial-lateral direction, such that number of participants who detected the 2 μm feature above chance levels with the index finger in medial-lateral sweeps was almost the same the number who performed above chance using their little fingers in the anterior-posterior direction.

### Limitations and future directions

Selecting stimulus height was a challenge in the present study. We used relatively large features (all above previously reported sensory thresholds) because we felt the additional difficulty of single-shot detection would make the task more difficult. Although this seemed to be true to some degree, shorter features would have allowed us to better estimate the true detection threshold and would have yielded more error trials which could have highlighted more important features of the movement strategy. We also did not give participants feedback on whether their selections were correct. Though we purposely did not allow for learning in this study, with feedback participants may have been able to improve their performance by further adjusting their behavioral strategies.

An interesting theme for future research is to investigate whether and how movement parameters change as tactile acuity declines with aging. Kennedy and colleagues (Kennedy et al., 2011) recorded the time it took participants to perform a graded series of detection procedures as a potential performance measure but did not quantify the way people moved their fingers for detection. One of the most interesting findings from that study is that tactile detection thresholds were unchanged across decades of lifespan, even as receptor density decreased.

Although tactile detection thresholds did not show a significant change in their cross-sectional study, the time to complete a graded detection test increased notably in older participants. The combination of these findings suggests that individuals develop movement strategies over lifetime to compensate for denervation. By limiting our participants to a single contact with each surface it would be easier to link a behavioral strategy with detection success.

## Acknowledgements

The authors thank Tim Goldhawk and Todd Simpson at the Western Nanofabrication facility and Bryan Dalrymple and Frank Van Sas at the Western Physics & Astronomy machine shop for manufacturing and preparing the stimuli used in these experiments.

## Author Roles

S.R. and J.A.P. conceived and designed research; S.R. and W.R.P. performed experiments; S.R. and R.Z.Z. analyzed data; S.R. prepared figures; S.R. and R.Z.Z. drafted manuscript; S.R. and J.A.P. edited and revised manuscript; S.R. W.R.P., R.Z.Z. and J.A.P. approved the final version of the manuscript.

